# One Section, Two Worlds: Single-Cell Integration of MALDI-MSI and Spatial Transcriptomics on the Same Single Tissue Section

**DOI:** 10.1101/2025.07.16.665110

**Authors:** Tim F.E. Hendriks, Gert B. Eijkel, Theodoros Visvikis, Benjamin Balluff, Ron M.A. Heeren, Eva Cuypers

## Abstract

Understanding tissue complexity requires spatially resolved multiomic data at single-cell resolution. Here, we present a workflow that integrates high-resolution matrix-assisted laser desorption ionization mass spectrometry imaging (MALDI-MSI) with Xenium spatial transcriptomics (SPT) on a single tissue section. This one-section strategy ensures exact spatial correspondence between metabolic and transcriptomic features, avoiding the misalignment issues of serial sections, where even minor offsets can result in sampling different cells. We validate compatibility of MALDI-MSI with downstream SPT, preserving transcriptomic quality despite semi-destructive ionization. Using mouse brain and human glioblastoma tissues, we achieve pixel-perfect modality coregistration, enabling per-cell MALDI spectra extraction aligned with gene expression. Integrated clustering reveals enhanced cell-type resolution and identifies metabolic heterogeneity within transcriptionally defined populations. This enables a direct and precise correlation between what a cell is doing and its biochemical state, providing a more holistic and accurate picture of cellular function, heterogeneity, and interaction in health and disease. Our workflow provides a scalable path to multiomic atlases of disease and development, advancing both data integration and translational research.

---

Understanding the spatial distribution of biomolecules and their interactions is key for unravelling the complexities of biological systems, particularly in complex systems such as brain ^1^. Modalities such as matrix-assisted laser desorption ionization-mass spectrometry imaging (MALDI-MSI) and spatial transcriptomics (SPT) have revolutionized spatial biology by enabling the visualization of metabolic and transcriptomic information directly within tissue sections^2, 3^. However, the integration of these modalities at a single cell level within the same tissue section remains a challenging yet promising area, offering unique insights into the molecular and cellular landscape of tissues.

MALDI-MSI enables the spatial detection of a wide range of molecular species, including lipids, metabolites, and proteins at a single cell level ^4, 5^. Although MALDI-MSI is not inherently suitable for the detection of larger RNA panels due to the technical challenges associated with ionizing and detecting nucleic acids using MALDI-MSI^6, 7^. RNA molecules are chemically distinct and more prone to degradation during sample preparation, and their relatively low abundance and complexity of ionization profiles further limit their detection with MALDI ^8^. SPT, particularly with cutting-edge platforms like Xenium, complements this approach by providing single-cell resolution maps of RNA expressions ^9^. The integration of these two modalities provides a comprehensive view of the molecular and transcriptomic states of individual cells, creating a deeper understanding of the interplay between molecular and genomic layers ^10^. One of the advantages of integrating MALDI-MSI and SPT on the same tissue section lies in the removal of serial-section variability. When both modalities are performed on the same section, the spatial registration of data from both modalities is essentially the same, reducing artifacts introduced by tissue heterogeneity and sectioning. This approach also ensures that molecular and transcriptomic data originate from the exact same cells and microenvironmental niches, preserving spatial fidelity and increasing confidence in biological inferences. Nevertheless, these clear single section multi-omics advantages the potential impact of MALDI-MSI on subsequent spatial transcriptomics quality and performance has not been systematically evaluated. Research by Godfrey *et al*, has shown that combining DESI-MSI and Visium allow for understanding the relationship between gene expression and the metabolic phenotype of cells in the complex tumour microenvironment ^11^. It needs to be mentioned that Visium lacks true single-cell resolution, it instead captures transcriptomes from spatially defined spots encompassing multiple cells. Consequently, Visium does not enable true single cell spatial multiomics. Therefore, we have chosen Xenium for our spatial single cell approach.

Achieving single-cell resolution is essential for understanding tissue architecture and cellular dynamics in complex tissues. While previous studies have already integrated spatial transcriptomics and molecular imaging, they often lack the resolution required to resolve individual cells, instead providing data averaged over larger regions or tissue compartments ^11-13^. This hinders accurate mapping of molecular features to specific cell types. In contrast, our approach enables direct correlation between transcriptomic identity and molecular phenotype at the individual cell level. This capability is critical for uncovering spatial relationships, identifying functional cell states, and detecting previously unresolved subpopulations ^14^. By preserving both the spatial and cellular integrity, single-section integration of MALDI-MSI and spatial transcriptomics enhances the reliability of downstream analyses, such as cell type-specific molecular profiling and multimodal machine learning, which are otherwise confounded by tissue misalignment and variability across serial sections.

In this study, we assess the effect of MALDI-MSI prior to Xenium SPT and demonstrate the feasibility of integrating 5 μm spatial resolution MALDI-MSI measurements and the single-cell Xenium spatial transcriptomics platform on the same tissue section. By leveraging the complementary strengths of these technologies, we established a workflow that combines single-cell MALDI-MSI with single-cell SPT data. To our knowledge, this is the first demonstrations of true single-section multimodal spatial analysis at cellular resolution, representing a significant technical advancement in spatial biology. As a proof of concept, we applied this approach to sagittal mouse brain sections, to provide a robust model to explore the integration of MALDI-MSI and SPT. The effect of MALDI-MSI on tissue integrity and the quality of spatial transcriptomics were evaluated to assess the impact of one technology on the other. Using human glioblastoma (GBM) tissue we established methods to spatially align and integrate data from MALDI-MSI and Xenium at single-cell resolution. We show the ability to obtain cell-specific MALDI-MSI spectra, linking molecular profiles to defined cell types. This work establishes a foundational framework for future studies that aim to link metabolic and transcriptomic data within single tissue sections on a per-cell basis providing powerful new insights into health and disease.

## Results

### Assess effects of prior MALDI-MSI on Spatial Transcriptomics Tissue Integrity and Data Quality

We first performed a series of validation experiments on adult mouse brain tissue to establish the feasibility of sequentially combining high-resolution MALDI-MSI with SPT. These preliminary studies served two purposes: to verify that the MALDI laser ablation and matrix application would not irreversibly damage tissue morphology, and to confirm that the semi-destructive MALDI step would not compromise downstream RNA capture or hybridization performance. Fresh-frozen mouse brain sections were mounted on Xenium-compatible slides, sublimated with 2,5-DHB matrix, and subjected to MALDI-MSI at 5 μm pixel size. Post-MSI, the same section underwent the Xenium SPT workflow. Cell boundaries were defined using Xenium Ranger. We observed that the MALDI-laser ablation produced a regular grid of micron-scale fluorescent spots within the intracellular matrix (fig. 1-A), corresponding precisely to the MALDI-MSI pixel array. These ablation marks did not obscure nuclear outlines, nor did they interfere with subsequent cell segmentation or identification. We directly compared counts in MALDI-MSI prior to SPT versus no MALDI-MSI prior to SPT (control) sections of mouse brain to quantify the impact of the MALDI laser on the transcript counts. Fig. 1-B, shows the transcript counts per cell within the defined cell boundaries per condition. In fig. 1-C, we aggregated transcript counts across five anatomically matched 1 mm^2^ regions in MALDI-MSI versus control sections. Boxplots of total RNA counts per cell and detected transcript count per mm^2^ reveal modest but statistically significant reductions in both RNA yield and transcript detection following MALDI-MSI. Although MALDI measured regions exhibited moderate statistically significant reductions in total RNA counts (-30%, p<0.005) and detected transcripts per cell (-20%, p<0.05) compared to control tissues, more than 90 % of cells in ablated zones still surpassed 250 detected transcripts. Despite this decrease, overall transcriptome complexity remained high, confirming that the semi-destructive MALDI step imposes only minor impacts on the spatial transcriptome.

**Figure 1:**
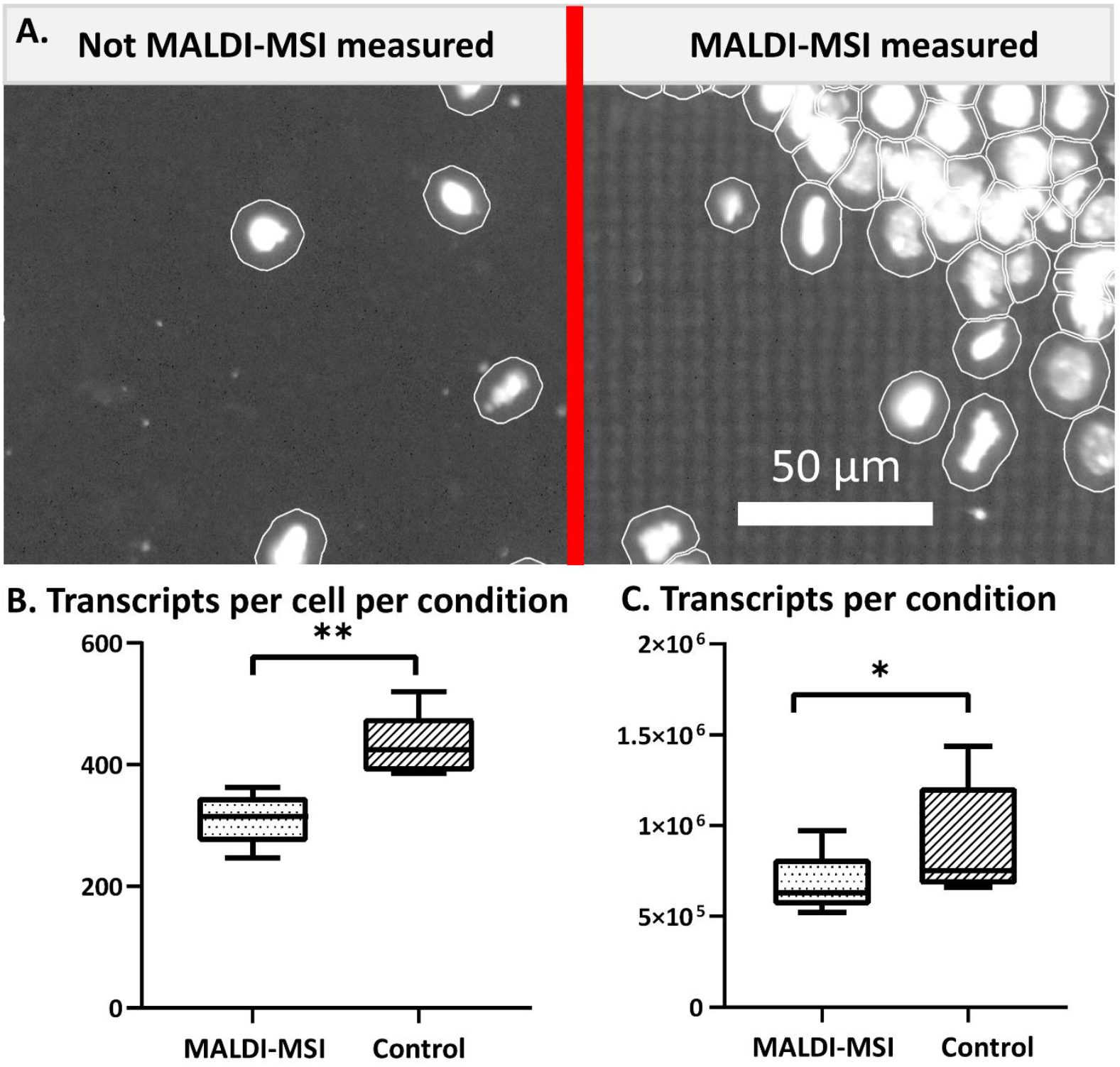
Impact of MALDI-MSI on tissue and spatial transcriptomics data quality. **(A)** DAPI-based cell segmentation control versus MALDI-MSI regions. Maximum-intensity projections of DAPI fluorescence in control (left) and MALDI-MSI (right) measured mouse brain. White outlines indicate cell boundaries defined by DAPI segmentation with a 5 μm radial expansion. The MALDI spot grid appears as a regular array of fluorescent microdots on the right but does not interfere with nuclear detection or boundary detection. Scale bar: 50 μm. Boxplots compare **(B)** total transcript counts per cell and **(C)** number of detected transcripts per area (mm^2^) between MALDI-MSI measured and non-MALDI-MSI measured (Control) mouse brain sections (≥40,000 cells per section). For the transcripts per area, 5 anatomical similar regions of 1mm^2^ were analysed on both tissues. Statistical significance was assessed by paired student T-test (** p < 0.005; * p < 0.05).

### Integration of MALDI-MSI with Xenium Spatial Transcriptomics

The integration of MALDI-MSI and SPT provides comprehensive multimodal insights into tissue biology at single-cell resolution. Here, we present the detailed overview of the computational workflow designed for integrating these two distinct, but complementary datasets obtained from the same tissue section (fig. 2-A). Firstly, MALDI-MSI and SPT measurements were sequentially performed on the same tissue section. The section was stained with a panel of fluorescent markers to enable cell segmentation: DAPI for nuclear identification, ATP1A1/CD45/E-Cadherin to define cell boundaries, 18S ribosomal RNA for interior RNA distribution, and αSMA/Vimentin for interior protein architecture. Following data acquisition, computational integration of both datasets is performed using an in-house MATLAB-based software platform we named ‘ESCDAT’ (sup. fig. 1). ESCDAT enables data handling and seamless spatial alignment between MALDI-MSI ion maps and spatial transcriptomics fluorescence images. Within ESCDAT, MALDI-MSI data is first binned into a lower resolution dataset (using a bin width of 1 Dalton), significantly enhancing computational efficiency and facilitating rapid preliminary analyses. Simultaneously, a fluorescence image from the spatial transcriptomics measurement is selected as the spatial reference for coregistration (fig. 2-B). Figure 2-B. illustrates the practical approach to accurately overlay MALDI-MSI data with corresponding spatial transcriptomics data. In panel *I*, a representative *m/z* channel from the MALDI-MSI analysis is visualized, providing spatial distribution of the selected molecule across the entire tissue section. Panel *II* shows a fluorescence image of the same tissue section, displaying cellular and molecular structures through labelled fluorescence markers which serve as reference data for integration. A focused examination of the region indicated by the red boxes in panels *I* and *II* is presented in panels *III* through *VI*. Panel *III* demonstrates a zoomed-in MALDI-MSI image revealing a sharp corner of a deliberately created fiducial marker region. The corresponding fluorescence image (panel *IV*) distinctly shows tissue ablation dots created by the MALDI laser for each pixel, which were strategically utilized as fiducial markers for precise pixel coregistration. In panel *V*, the overlay of MALDI-MSI sampling points (red circles) onto the fluorescence image demonstrates accurate spatial alignment achieved by the coregistration method. Panel *VI* further highlights this alignment by presenting the MALDI-MSI data combined with cell boundary segmentation derived from SPT. The alignment of these modalities leveraged fiducial markers identifiable in both MALDI-MSI and fluorescence images. This step ensured precise spatial coregistration, essential for accurate downstream biological interpretation. Following spatial alignment, cell boundary coordinates obtained via Xenium Ranger using the cell segmentation staining were imported into ESCDAT, allowing for extraction of high-resolution per-cell *m/z* spectra from the MALDI-MSI data. The resulting integrated multimodal dataset, combined high-resolution mass spectra and spatially resolved gene expression profiles, was then analysed using Seurat v5. This integrated visualization clearly demonstrates how molecular data from MALDI-MSI align spatially with single-cell transcriptomic data, thereby enabling detailed multiomic cellular profiling. Additional observations of interest include the precision and spatial resolution attainable through this method, confirming the potential in correlating distinct metabolic profiles identified by MALDI-MSI directly with specific cell types and their transcriptional states.

**Figure 2.**
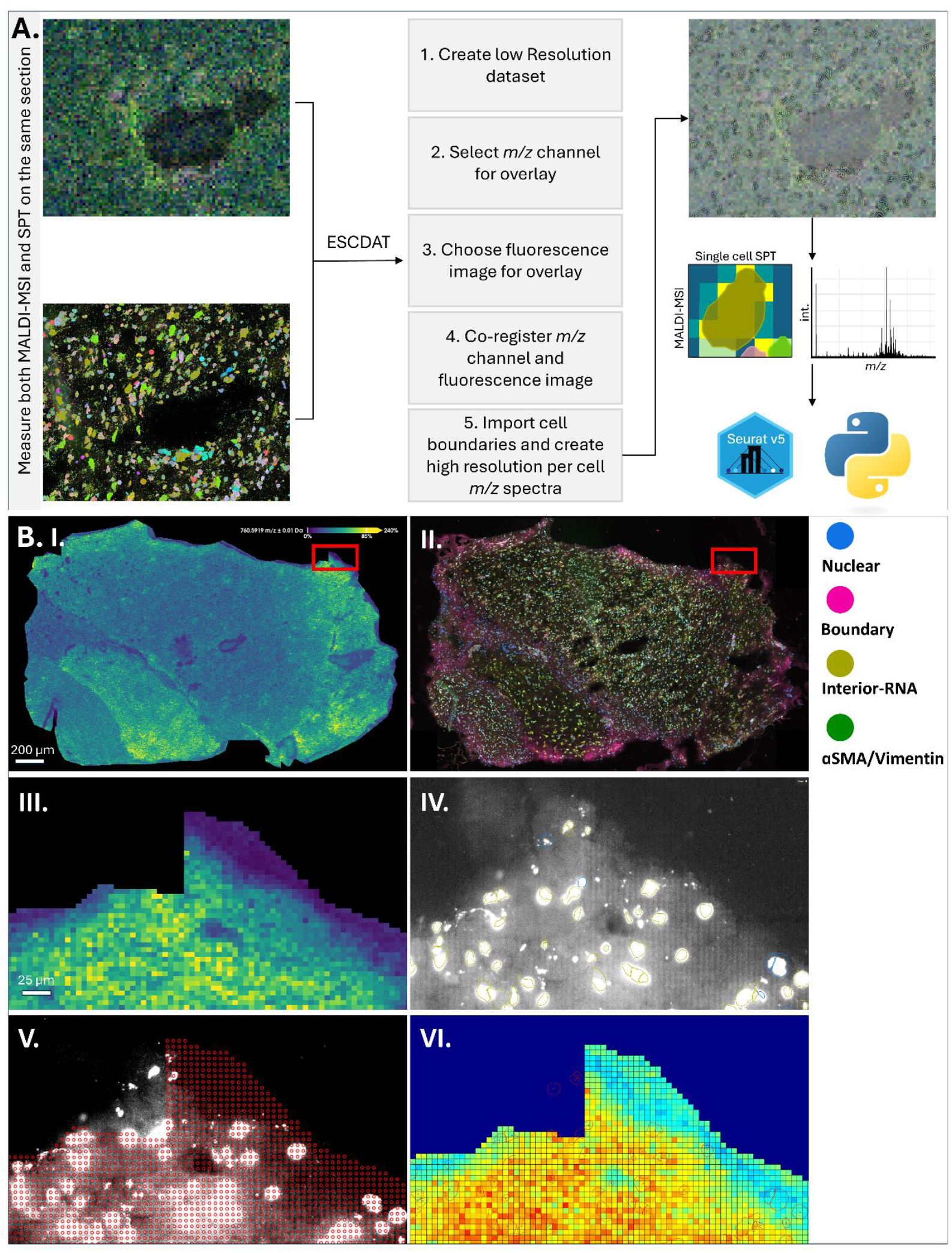
Overview of overlaying and combining MALDI-MSI and spatial transcriptomics data. **(A)** Shows the schematic workflow used using ESCDAT. First MALDI-MSI (TimsTOF fleX) and spatial transcriptomics (Xenium) are sequentially conducted on the same tissue section. Afterwards, using ESCDAT, a low mass resolution dataset is spatially aligned with the spatial transcriptomics data. Using fiducial markers and visible laser spots in the tissue generated via the MALDI laser. Using the cell boundaries from the cell segmentation staining, pixels per cell are averaged and a high-resolution *m/z* spectrum is created per cell. These data modalities are afterwards combined, and downstream analysis are performed using Seurat V5 and Python. **(B)** Visualization of coregistration MALDI-MSI with spatial transcriptomics. *(I)* A representative m/z ion map from MALDI-MSI showing the spatial distribution of a selected molecule across the tissue. *(II)* Fluorescence image of the same section with labeled markers as reference. *(III-IV)* Zoom into the red-boxed region: *(III)* MALDI-MSI view of a sharp fiducial marker corner; *(IV)* matching fluorescence view with the laser-ablation dots used as fiducials. *(V)* Overlay of MALDI sampling points (red circles) on the fluorescence image after coregistration. *(VI)* Final overlay of MALDI-MSI data withcell-boundary segmentation from spatial transcriptomics, demonstrating precise multimodal alignment.

### Cell identification and Cell-specific MALDI-MSI Spectra

We profiled 8,318 cells across the tissue that overlapped between MALDI-MSI and SPT, after applying filters to remove low quality cells (qv < 20). Initial dataset quality was assessed by examining total RNA counts (nCount_RNA) and detected transcript numbers (nFeature_RNA) across all predicted cell-type clusters (sup. fig 2). The similar shapes and ranges of these distributions confirm consistent capture efficiency and sequencing depth amongst the cell types. Accurate cell-type identification was performed based on spatial transcriptomic profiles using Seurat v5 and log-normalization, leveraging predefined marker genes for distinct cell populations (sup. table 1) ^15-27^. The cell identifications were set as a ground-truth for the cell identities throughout. These marker genes, selected based on prior biological knowledge, demonstrated high specificity and robust expression patterns within targeted cell types (sup. fig. 3). In every cell type panel, the distribution of scores peaks sharply in the matching cell-type group and remains low in all others. For example, the astrocyte signature has its highest values exclusively in cells predicted as astrocytes, and the endothelial cell signature peaks only in endothelial-predicted cells. Minor low-level signals in non-target groups reflect baseline expression and technical background. Further validation at the single-gene level confirms that each marker’s normalized expression is tightly confined to its predicted lineage. Analysis of individual marker-gene expression distributions (sup. fig. 4) confirms that each gene is predominantly expressed in its assigned cell-type cluster, with negligible off-target expression. For instance, NRP1 and PECAM1 localize exclusively to endothelial cells, and MOG and CAPN3 peaks are restricted to oligodendrocytes. These plots validate both our marker lists and the prediction strategy: each marker panel cleanly identifies its intended lineage with minimal crosstalk, further substantiating the accuracy of our annotations. To further resolve the transcriptional heterogeneity underlying our spatially defined cell-type assignments, we used Seurat to identify the top ten positively enriched genes per predicted cell type (ranked by average log_2_-fold change). We subsequently visualized the normalized expression of these 100 marker genes across all cells in a heatmap (sup. fig. 5), with cells ordered by their predicted cell type. Across all clusters, marker-gene expression is sharply confined to its cognate cell-type segment, with minimal off-target signals. Similar analysis was performed for the MALDI-MSI markers. Using Seurat on the TIC-normalized MALDI assay, we generated a heatmap of the top 100 *m/z* features based on log-fold-change across all cell types for all cells (sup. fig. 6). Cells remain ordered by their predicted cell type. Unlike the binary on/off patterns seen in transcript data, MALDI-MSI data exhibits graded enrichment: certain *m/z* values are elevated in one cell type yet maintain low-level expression elsewhere. These subtler, non-binary distributions reflect the shared metabolic milieu of neighbouring cell types yet still show clear molecular signatures that can complement the RNA-based annotations.

### Combined Molecular Profiling of Single Cell Data

Next, we investigated whether the integration of SPT and MALDI-MSI data significantly enhances the resolution of single-cell clustering compared to either modality alone. Before dimensionality reduction, both datasets were centered, scaled, and the top 30 PCs were retained for UMAP construction. The individual PCA and UMAP plots for RNA and MALDI-MSI data revealed notable differences in cell-type resolution. RNA PCA and UMAP (sup. fig. 7-A) reveals clear clustering, especially the RNA UMAP demonstrated the capability of spatial transcriptomics data to differentiate and delineate distinct cell populations. Clusters such as oligodendrocytes, excitatory, and inhibitory neurons, endothelial cells, and glial stem cells (GSCs) are clearly separated. The MALDI-MSI PCA and UMAP (sup. fig. 7-B) reveal limited clustering, indicating that MALDI-MSI alone might insufficiently discriminate between the different neuronal cell types. Although very subtle structural groupings are observable the specific cell identities are not clearly separated. This comparative analysis underscores the higher cell-type discriminatory power of spatial transcriptomics data when compared to MALDI-MSI alone. However, when combining the RNA and MALDI-MSI data per cell through multimodal neighbour analysis, the resulting UMAP visualization (fig. 3-A) shows new subclusters. Distinct cell populations, including oligodendrocytes, astrocytes, endothelial cells, TAMs, and neuronal subpopulations, were clearly delineated. The integrated modalities can capture complementary molecular information, thus greatly refining the spatial resolution and biological interpretation compared to either modality alone. It also showed the value and necessity of integrating these complementary data modalities to enhance biological relevance. While RNA markers provide clear cell-type identification, MALDI-MSI data enriched the molecular profiling, particularly evident from the combined UMAP clusters. The modality contribution analysis (fig. 3-B) reveals distinct patterns in the reliance on RNA versus MALDI-MSI data for clustering per cell type. Certain cell types, including astrocytes, neuronal populations, endothelial cells, GSCs and MES cells showed balanced modality contributions (close to a 50/50 ratio), suggesting equally important roles of transcriptomic and metabolic signatures for clustering. In contrast, other populations, such as oligodendrocytes, OPCs, TAMs and T-cells predominantly relied on RNA for accurate clustering. The increase of information by integrating SPT and MALDI-MSI is supported by visualizing all transcripts and all *m/z* values (sup. fig. 8) in a correlation heatmap. Here we saw several coherent clusters that show either positive or negative correlation. By systematically exploring the associations between top RNA marker genes characteristic of specific cell types and top MALDI-MSI ions (fig. 3-C). Here we notably saw correlations emerge between distinct cell-type markers and particular ions, suggesting functional and metabolic specificity for these cell populations. This correlation heatmap further suggests precise associations between specific transcripts and molecular ions, offering insights into cell-type-specific biological processes or metabolic pathways. These findings highlight variability in the molecular information content contributed by each modality across cell types and emphasize the complementary strengths of integrating both RNA and MALDI-MSI data to achieve additional biological information and understanding.

**Figure 3.**
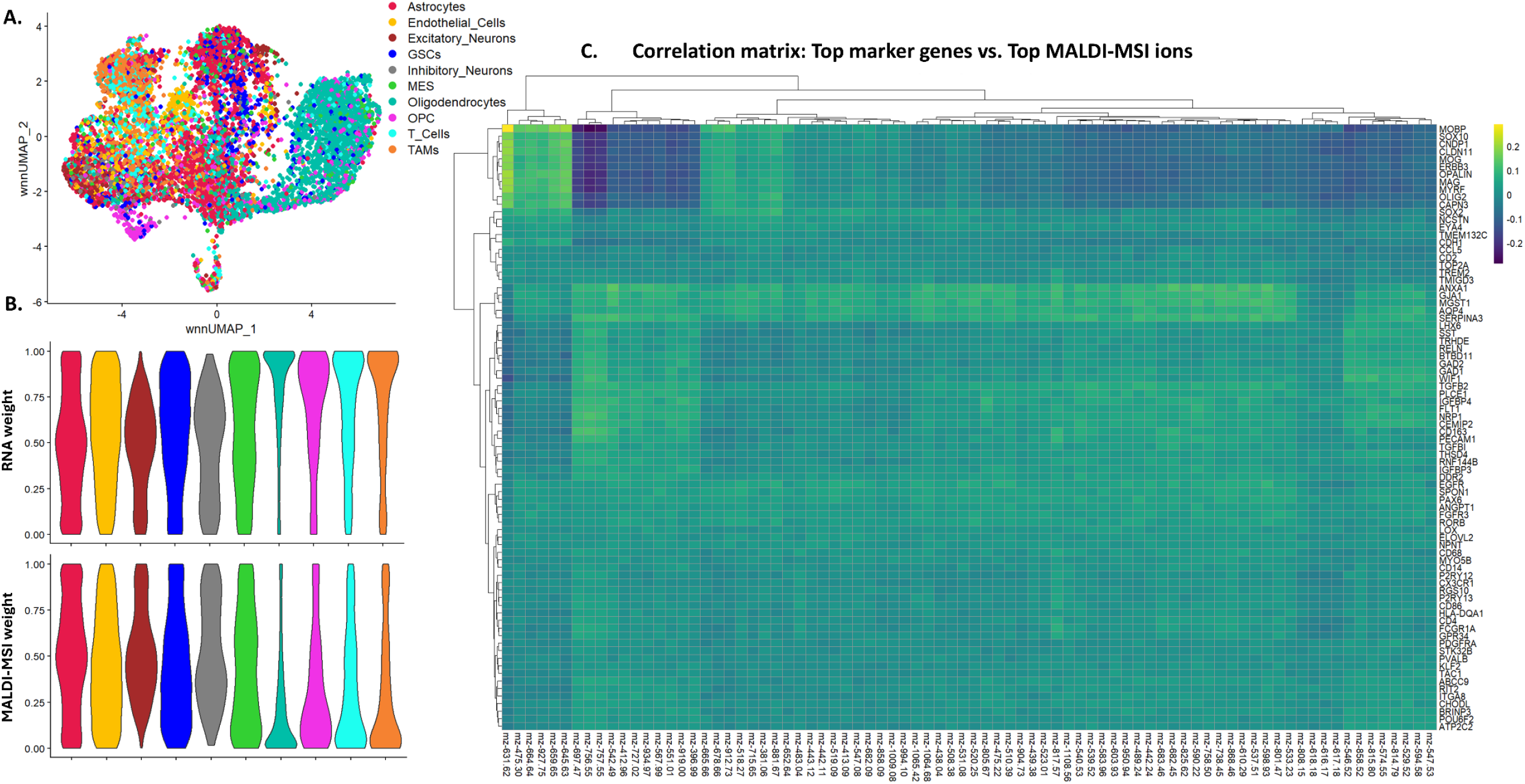
Integrated multimodal single-cell molecular profiling reveals distinct cell clusters. **(A)** Combined UMAP visualization of integrated spatial transcriptomics (RNA) and MALDI-MSI data. Distinct cell-type clusters, labelled according to RNA-based ground-truth cell identities, are clearly distinguishable when both molecular modalities are combined. **(B)** Violin plots showing the contribution of each modality (RNA vs. MALDI-MSI) to cell-type clustering. Notably, some cell populations, such as astrocytes, endothelial cells, excitatory and inhibitory neurons, GSCs and MES cells, exhibit balanced contributions (∼50% RNA and 50% MALDI-MSI) to the UMAP. whereas oligodendrocytes, OPCs, T-cells and TAMs predominantly rely on RNA data, indicating variability in modality-specific information content across cell types. **(C)** Correlation heatmap between top 80 RNA marker genes and top 80 MALDI-MSI ions. Correlation patterns suggest significant interdependencies between RNA expression levels and molecular ions, hinting at underlying biological or metabolic pathways specific to certain cell types.

## Discussion

In this study, we established and demonstrate a workflow for integrating high-resolution MALDI-MSI with Xenium single cell spatial transcriptomics on one single tissue section, overcoming the limitations of prior approaches that relied on adjacent sections or lower-resolution coregistration ^11, 13, 28^. This approach is motivated by the need to achieve a comprehensive, spatially precise, and cell-specific understanding of tissue biology. This cannot be reliably achieved using either modality alone or by analysing adjacent sections. First, validating our approach on mouse brain before applying it to human glioblastoma. We found that sublimation of 2,5-DHB and laser ablation at a 5 μm pixel pitch produced a characteristic fluorescent grid of ablation spots, yet nuclear morphology and cell segmentation remained entirely intact. Importantly, we performed MALDI-MSI directly on Xenium slides, without requiring conventional conductive ITO-coated substrates, demonstrating that high-resolution MALDI imaging is feasible on transcriptomics compatible surfaces using TimsTof. This removes a key compatibility barrier between platforms, simplifies multimodal sample preparation, and enables broader adoption of integrated workflows. While the integration was successful, we observed moderate reductions in transcript counts following MALDI-MSI without any significant effect on cell type identifications ^29^. Indicates that the transcriptomic data remains biologically informative, supporting reliable cell-type classification and downstream analyses, making the trade-off acceptable for integrated multimodal workflows.

Using the regular grid of MALD-MSI laser ablation spots, ESCDAT enabled pixel-perfect alignment of MSI ion maps with fluorescence-based cell segmentation, allowing per-cell mass spectra to be paired with spatial transcriptomes. While single cell RNA markers formed discrete, lineage-specific clusters, MALDI-MSI features showed graded biochemical enrichments. Cell-type specific masses were markedly elevated in their respective types but also showed low-level expression elsewhere, reflecting shared metabolic contexts^31^. Dimensionality-reduction embeddings further highlight this complementarity: RNA-only UMAPs resolve all ten canonical cell types into well separated islands, whereas MALDI-MSI UMAPs shows substantial overlap among cell identities. Yet, when integrated, the two modalities sharpen cluster boundaries and can increase confidence in cell-type assignments. Correlation heatmaps linking gene expression to m/z features revealed that specific cell types exhibit distinct combinations of transcripts and molecular ions, effectively defining unique cellular ‘metabotypes’. These gene-metabolite associations suggest that transcriptomic identity is tightly coupled to underlying biochemical states, enabling functional characterization of cells beyond RNA alone and suggests candidate gene-metabolite regulatory circuits for further investigation.

Beyond technical validation, these multimodal maps revealed new biological insights. The combined MALDI signals with RNA-defined clusters point to metabolic heterogeneity even among transcriptionally similar cells, hinting at subpopulations within the section. Unlike traditional 2D or 3D cultures, our single tissue-based workflow maintains cells in their native spatial and microenvironmental context, enabling the study of molecular and transcriptomic phenotypes as they exist *in situ* ^32^. This allows not only for more biologically relevant single cell profiling but also for the analysis of cell-to-cell interactions, by examining how neighbouring cells with distinct molecular identities influence one another through spatial proximity. Looking forward, expanding our workflow to include targeted lipidomic, protein markers, and advanced statistical integration frameworks will further extend the understanding of cellular states in health and disease. As transcriptomic platforms continue to grow in gene coverage, including full-transcriptome panels, our approach enables even more refined classification. By linking gene expression to cell-type-specific metabolic or signalling profiles, this integration enhances our ability to dissect disease mechanisms, identify biomarkers or drug targets, and distinguish cell states that are transcriptionally similar but functionally distinct. It also aligns with the goals of atlas-scale efforts, such as the Human Cell Atlas, by enabling the generation of spatially resolved, multiomic reference maps that preserve both cellular identity and microenvironmental context ^33, 34^. Ultimately, this integrated MALDI-MSI/SPT platform lays the groundwork for comprehensive, single-cell resolved multiomic atlases.

To enable broader adoption, key steps in our workflow such as matrix deposition, imaging, and registration are already compatible with high-throughput instrumentation. Our integration software (ESCDAT) is designed to be modular and will be further developed for other modalities and scalable deployment, such as multiple tissue sections. Compared with existing multimodal platforms, our method uniquely achieves subcellular coregistration of lipidomic and transcriptomic data without sacrificing either modality’s resolution. By generating truly cell-specific mass spectra that directly correspond to Xenium derived transcriptomes, our workflow sets a new standard for comprehensive multiomic tissue mapping, enabling more precise investigation of metabolic reprogramming in tumour microenvironments and delivering a superior tool for both fundamental research and translational pathology.

## Materials and methods

### Sample preparation – MALDI-MSI

Ethical approval was obtained from the UZ/KU Leuven Ethical Review Committee of University Hospitals Leuven (Gasthuisberg, Leuven, Belgium) with reference number S60290 and the project was registered under the UZ Leuven Tissue Bank. Fresh frozen mouse brain (n=3) and fresh frozen human GBM tumour tissue were used, where the latter was embedded in 10% gelatin 2% CMC (SigmaAldrich). Afterwards tissues were cryo-sectioned at 10 μm sections using a cryostat (CM1850, Leica Biosystems) according to Protocol CG000579 published by 10X Genomics. The sections were stored at −80°C until further analysis. Upon retrieval, they were immediately transferred to boxes filled with hygroscopic desiccant beads and subsequently vacuum-dried in a desiccation chamber for 20 minutes. Before MALD-MSI experiments, sublimation of 50 mg of 2,5-dihydroxybenzoic acid (Sigma-Aldrich) matrix at 160°C for 160 seconds was performed using an HTX Sublimator (HTX Technologies, USA).

### MALDI-MSI analysis

Whilst setting up the imaging-experiment, to create fiducial markers, several sharp corners and features on the tissue were created during the ROI selection (fig 2. B-III). MALDI-MSI analysis was performed on a timsTOF flex (Bruker Daltonics GmbH, Germany). Spectra were obtained in positive ion mode with a mass range of *m/z* 250 – 1200 with a pixel size of 5x5 μm and laser spot size of approximately 4 μm. The laser frequency was set to 1 kHz, and 25 shots were accumulated at each pixel. The MALDI laser consists of a Nd:YAG 355 nm SmartBeam 3D laser (Ekspla, Lithuania). Time-of-flight calibration was performed before imaging experiments using red phosphorus. Transfer settings were 200 V peak-to-peak (Vpp; funnel 1 RF), 350 Vpp (funnel 2 RF), and 450 Vpp (multipole RF). Focus pre-time-of-flight (TOF) transfer time was set at 100 μs. The quadrupole ion energy was 5.0 eV with a low mass of *m/z* 200. Collision cell energy was 10.0 eV with collision RF at 1500 Vpp.

### Sample preparation – Spatial Transcriptomics

After MALDI-MSI, to remove any access 2,5-DHB, slides were washed for two times 1 minute in 100% EtOH (BioSolve). Slides were then dried under a steady N_2_ stream. The Xenium slides were then fixed and permeabilized following Protocol CG000581. In addition to this protocol, slides were fully submerged in PBS at 37°C for 2 minutes. Afterwards, any gelatin covering the Xenium slide fiducials was manually removed using a tissue (KimTech wipes, Sigma Aldrich). Probe hybridization, ligation, and amplification were carried out using the pre-designed Mouse Brain and Human Brain reagent in accordance with Protocol CG000749. Reagents for the Xenium measurements were bought according to Protocol CG000601. Finally, the processed slides were loaded into the Xenium Analyzer as per Protocol CG000584. After the Xenium analysis, raw FASTQ files were analysed using Xenium Ranger (v2.0.0.10) using the Xenium multimodal cell segmentation method. Data was then visualized in Xenium Explorer (v3.2.0) and processed using the Seurat R package (v5.3.0).

### Integration of MALDI-MSI and spatial transcriptomics

Single cells detected via the Xenium multimodal cell segmentation were converted to a GeoJSON format containing the cell-identifier, cell cluster, individual transcript count and spatial coordinates using our custom python script. Afterwards, Tims-TOF MALDI-MSI and Xenium spatial transcriptomics datasets were aligned using our custom co-registration software (Version P1 04-06-2025) compiled in MATLAB (v2022b). First, the Bruker Tims-TOF binary (.tsf) files were imported via Python, and a low-resolution 1 Da mass-spectrometry image was generated. We then created either a total ion current (TIC) image or a single-mass-channel image and matched it to the fluorescence microscope image produced during Xenium’s multi-modal cell-segmentation staining. MALDI-laser ablation spots are readily visible in the fluorescence channel, providing precise markers for near-pixel-perfect alignment. Tissue fiducials placed before MALDI-MSI were co-localized with these laser spots to match at least four corresponding registration points. A piecewise-linear (PWL) transformation was then computed, producing a transformation matrix that links coordinates across modalities. Finally, we applied this matrix to the GeoJSON file exported from the Xenium run containing cell IDs, spatial coordinates, and transcript counts to extract high-resolution mass spectra for each cell-id within the combined coordinate frame. To account for partial-volume effects and varying cell-pixel overlap, we weighted each MALDI pixel’s intensity by the fraction of its area overlapping the cell segmentation mask, as demonstrated before ^35^.

### Data Processing and Cell Type Identification

Xenium data was processed using the Seurat R package (v5.3.0). Filtering of cells excluded all low-quality cells with a quality value (qv) of < 20. Also, any empty cells with an nCount lower than 0 were removed. Further analysis was only conducted on common cells between the MALDI-MSI and Xenium measurement. The MALDI-MSI data was imported as a new assay in the Xenium Seurat object. Data for both modalities were normalized and scaled. Afterwards, PCA was conducted on both for the optimal number of PCs, found via elbow plots and PC expression heatmaps. Dimensionality and visualization were performed using UMAP. Any clusters with overall low nCount were removed, and PCA and UMAP visualization were performed again on the filtered object. Cell identifications in the clusters were characterized and visualized using expressed marker genes from the Human Brain panel.

### Differential Marker Analysis Per Cell Type

To identify both transcriptomic and *m/z* features that are distinguishable for each predicted cell type, we performed cluster-wise differential testing using Seurat v5. First, for RNA markers, the RNA expression values were log normalized. We ran a *FindAllMarkers()* test which employs a Wilcoxon rank-sum test and controls the false discovery rate via Bonferroni correction. From the resulting table, we selected the top ten genes per cluster ranked by average log_2_-fold change and visualized in a grouped heatmap. For the *m/z* features, we used TIC-normalized MALDI data and performed similar tests. To jointly leverage transcriptomic and mass spectrometric modalities, we employed Seurat v5’s weighted nearest-neighbour (WNN) framework. First, we computed independent principal component analyses on the RNA and TIC-normalized MALDI assays, retaining the first 30 PCs from each modality. To build a shared neighbour graph weighted by both data types UMAP was performed. For feature-pair correlation analysis, we extracted the per-cell scaled matrices and then computed Pearson correlations between features. Hierarchical clustering was applied to both rows and columns.

## Supporting information

Supplemental figures MALDI-MSI and SPT

## Data availability

Raw and processed MALDI-MSI and Xenium spatial transcriptomics data from the mouse measurements, including Seurat objects, coregistration overlays, and associated analysis scripts, are available from the corresponding author upon reasonable request. Please note that glioblastoma data cannot be shared directly, as this falls outside the scope of the approved ethical consents. Access to this data requires additional institutional review and adherence to the original patient consent.

## Code availability

We used ESCDAT Version P1 04-06-2025 for the overlay in this manuscript. The software packages such as R, Seurat and Python are publicly available and well documented. The Xenium to GeoJSON package and MALDI-MSI – Xenium SPT software ‘ESCDAT’ underlying these analysis are deposited at GitHub (https://github.com/M4i-Imaging-Mass-Spectrometry/MALDI-MSI---Spatial-Transcriptomics-Overlay).

## Author information

### Corresponding Author

Eva Cuypers – Maastricht MultiModal Molecular Imaging Institute – M4i, Maastricht University, Maastricht 6229 ER, The Netherlands

### Authors

### Author contributions

T.F.E.H. contributed to the investigation, formal analysis, conceptualization, writing – review, and editing; G.B.E. contributed to software design and development, conceptualization and review; T.V. contributed to conceptualization, writing and review; B.B. contributed to software design and review; R.M.A.H. contributed to review and editing, funding acquisition; E.C. contributed to conceptualization, supervision, writing - review and editing, funding acquisition.

### Notes

The authors declare no competing financial interest.

### Supporting information

ST1 – Gene markers per cell type; SF1 – Overview GUI ESCDAT; SF2 – Gene counts and features per cell type; SF3 – Score distribution per cell type; SF4 – Score distribution of individual marker genes; SF5 – Heatmap top RNA markers; SF6 – Heatmap of top *m/z* markers; SF7 – Modality specific dimensionality reductions; SF8 – Global gene – *m/z* correlation heatmap.

## Acknowledgments

This research was supported by the LINK program funded through the Netherlands Organization for Scientific Research (NWO) and NWO-STEM (Project Number 19013 to E.C.). We also gratefully acknowledge the support of the FWO Research Foundation, Belgium (TBM T001919N) and the Interreg Flanders-Netherlands Molecular Brain Tumor Detector. The authors would like to thank Morgane Raoult, Henk Buermans and Najiba Mammadova (10X Genomics) for performing the mouse Xenium Spatial Transcriptomics experiment. Ruben Jacobs (Maastricht University) for performing the human glioblastoma Xenium Spatial Transcriptomics experiment and Ian G.M. Anthony (Maastricht University) for helping with the python code. The authors would also like to thank the Department of Neurosurgery at KU Leuven, especially Steven de Vleeschouwer, for providing the GBM sample.

## References

(1) Fangma, Y.; Liu, M.; Liao, J.; Chen, Z.; Zheng, Y. Dissecting the brain with spatially resolved multi-omics. J Pharm Anal 2023, 13 (7), 694–710. DOI: 10.1016/j.jpha.2023.04.003 From NLM PubMed-not-MEDLINE.

(2) Janesick, A.; Shelansky, R.; Gottscho, A. D.; Wagner, F.; Williams, S. R.; Rouault, M.; Beliakoff, G.; Morrison, C. A.; Oliveira, M. F.; Sicherman, J. T.; et al. High resolution mapping of the tumor microenvironment using integrated single-cell, spatial and in situ analysis. Nat Commun 2023, 14 (1), 8353. DOI: 10.1038/s41467-023-43458-x From NLM Medline.

(3) Mirzazadeh, R.; Andrusivova, Z.; Larsson, L.; Newton, P. T.; Galicia, L. A.; Abalo, X. M.; Avijgan, M.; Kvastad, L.; Denadai-Souza, A.; Stakenborg, N.; et al. Spatially resolved transcriptomic profiling of degraded and challenging fresh frozen samples. Nat Commun 2023, 14 (1), 509. DOI: 10.1038/s41467-023-36071-5 From NLM Medline.

(4) Sekera, E. R.; Rosas, L.; Holbrook, J. H.; Angeles-Lopez, Q. D.; Khaliullin, T.; Rojas, M.; Mora, A. L.; Hummon, A. B. Single Cell MALDI-MSI Analysis of Lipids and Proteins within a Replicative Senescence Fibroblast Model. J Am Soc Mass Spectrom 2024, 35 (12), 2815–2823. DOI: 10.1021/jasms.4c00095 From NLM Medline.

(5) McKinnon, J. C.; Milioli, H. H.; Purcell, C. A.; Chaffer, C. L.; Wadie, B.; Alexandrov, T.; Mitchell, T. W.; Ellis, S. R. Enhancing metabolite coverage in MALDI-MSI using laser postionisation (MALDI-2). Anal Methods 2023, 15 (34), 4311–4320. DOI: 10.1039/d3ay01046e From NLM Medline.

(6) Zhang, H.; Lu, K. H.; Ebbini, M.; Huang, P.; Lu, H.; Li, L. Mass spectrometry imaging for spatially resolved multi-omics molecular mapping. Npj Imaging 2024, 2 (1), 20. DOI: 10.1038/s44303-024-00025-3 From NLM PubMed-not-MEDLINE.

(7) Kyle A. Vanderschoot, J.P.P., Kelli A. Steineman, Christopher M. De Caro, Marie C. Heffern, Elizabeth K. Neumann. Spatial Multiomics Lipids and Gene expression using MALDI ISH MSI. BioRxiv 2024. DOI: 10.1101/2024.06.01.596997.

(8) Emanuelson, C.; Ankenbruck, N.; Deiters, A.; Yu, M. S. High-Throughput Amenable MALDI-MS Detection of RNA and DNA with On-Surface Analyte Enrichment Using Fluorous Partitioning. SLAS Discov 2021, 26 (1), 58–66. DOI: 10.1177/2472555220958391 From NLM Medline.

(9) Liu, Q.; Shen, C.; Dai, Y.; Tang, T.; Hou, C.; Yang, H.; Wang, Y.; Xu, J.; Lu, Y.; Wang, Y.; et al. Single-cell, single-nucleus and xenium-based spatial transcriptomics analyses reveal inflammatory activation and altered cell interactions in the hippocampus in mice with temporal lobe epilepsy. Biomark Res 2024, 12 (1), 103. DOI: 10.1186/s40364-024-00636-3 From NLM PubMed-not-MEDLINE.

(10) Zheng, P.; Zhang, N.; Ren, D.; Yu, C.; Zhao, B.; Zhang, Y. Integrated spatial transcriptome and metabolism study reveals metabolic heterogeneity in human injured brain. Cell Rep Med 2023, 4 (6), 101057. DOI: 10.1016/j.xcrm.2023.101057 From NLM Medline.

(11) Trevor M. Godfrey, Y.S., Wanqiu Zhang, Thao Tran, Nico Verbeeck, Nathan H. Patterson, Faith E. Jackobs, Chandandeep Nagi, Maheshwari Ramineni, Livia S. Eberlin. Integrating Ambient Mass Spectrometry Imaging and Spatial Transcriptomics on the Same Cancer Tissues to Identify Gene-Metabolite Correlations. BioRxiv 2024. DOI: 10.1101/2024.12.20.626670.

(12) Kreutzer, L.; Weber, P.; Heider, T.; Heikenwalder, M.; Riedl, T.; Baumeister, P.; Klauschen, F.; Belka, C.; Walch, A.; Zitzelsberger, H.; et al. Simultaneous metabolite MALDI-MSI, whole exome and transcriptome analysis from formalin-fixed paraffin-embedded tissue sections. Lab Invest 2022, 102 (12), 1400–1405. DOI: 10.1038/s41374-022-00829-0 From NLM Medline.

(13) Vicari, M.; Mirzazadeh, R.; Nilsson, A.; Shariatgorji, R.; Bjarterot, P.; Larsson, L.; Lee, H.; Nilsson, M.; Foyer, J.; Ekvall, M.; et al. Spatial multimodal analysis of transcriptomes and metabolomes in tissues. Nat Biotechnol 2024, 42 (7), 1046–1050. DOI: 10.1038/s41587-023-01937-y From NLM Medline.

(14) Huynh, K. L. A.; Tyc, K. M.; Matuck, B. F.; Easter, Q. T.; Pratapa, A.; Kumar, N. V.; Perez, P.; Kulchar, R.; Pranzatelli, T.; de Souza, D.; et al. Spatial Deconvolution of Cell Types and Cell States at Scale Utilizing TACIT. bioRxiv 2024. DOI: 10.1101/2024.05.31.596861 From NLM PubMed-not-MEDLINE.

(15) Li, J.; Wang, P.; Wang, L. Y.; Wu, Y.; Wang, J.; Yu, D.; Chen, Z.; Shi, H.; Yin, S. Redistribution of the astrocyte phenotypes in the medial vestibular nuclei after unilateral labyrinthectomy. Front Neurosci 2023, 17, 1146147. DOI: 10.3389/fnins.2023.1146147 From NLM PubMed-not-MEDLINE.

(16) Yang, Y.; Chu, L.; Zeng, Z.; Xu, S.; Yang, H.; Zhang, X.; Jia, J.; Long, N.; Hu, Y.; Liu, J. Four specific biomarkers associated with the progression of glioblastoma multiforme in older adults identified using weighted gene co-expression network analysis. Bioengineered 2021, 12 (1), 6643–6654. DOI: 10.1080/21655979.2021.1975980 From NLM Medline.

(17) Amin, M.; Mays, M.; Polston, D.; Flanagan, E. P.; Prayson, R.; Kunchok, A. Myelin oligodendrocyte glycoprotein (MOG) antibodies in a patient with glioblastoma: Red flags for false positivity. J Neuroimmunol 2021, 361, 577743. DOI: 10.1016/j.jneuroim.2021.577743 From NLM Medline.

(18) Ye, D.; Wang, Q.; Yang, Y.; Chen, B.; Zhang, F.; Wang, Z.; Luan, Z. Identifying Genes that Affect Differentiation of Human Neural Stem Cells and Myelination of Mature Oligodendrocytes. Cell Mol Neurobiol 2023, 43 (5), 2337–2358. DOI: 10.1007/s10571-022-01313-5 From NLM Medline.

(19) Hayashida, S.; Masaki, K.; Suzuki, S. O.; Yamasaki, R.; Watanabe, M.; Koyama, S.; Isobe, N.; Matsushita, T.; Takahashi, K.; Tabira, T.; et al. Distinct microglial and macrophage distribution patterns in the concentric and lamellar lesions in Balo’s disease and neuromyelitis optica spectrum disorders. Brain Pathol 2020, 30 (6), 1144–1157. DOI: 10.1111/bpa.12898 From NLM Medline.

(20) Mecca, C.; Giambanco, I.; Donato, R.; Arcuri, C. Microglia and Aging: The Role of the TREM2-DAP12 and CX3CL1-CX3CR1 Axes. Int J Mol Sci 2018, 19 (1). DOI: 10.3390/ijms19010318 From NLM Medline.

(21) Ravi, V. M.; Neidert, N.; Will, P.; Joseph, K.; Maier, J. P.; Kuckelhaus, J.; Vollmer, L.; Goeldner, J. M.; Behringer, S. P.; Scherer, F.; et al. T-cell dysfunction in the glioblastoma microenvironment is mediated by myeloid cells releasing interleukin-10. Nat Commun 2022, 13 (1), 925. DOI: 10.1038/s41467-022-28523-1 From NLM Medline.

(22) Lichtenberger, B. M.; Tan, P. K.; Niederleithner, H.; Ferrara, N.; Petzelbauer, P.; Sibilia, M. Autocrine VEGF signaling synergizes with EGFR in tumor cells to promote epithelial cancer development. Cell 2010, 140 (2), 268–279. DOI: 10.1016/j.cell.2009.12.046 From NLM Medline.

(23) Tasic, B.; Yao, Z.; Graybuck, L. T.; Smith, K. A.; Nguyen, T. N.; Bertagnolli, D.; Goldy, J.; Garren, E.; Economo, M. N.; Viswanathan, S.; et al. Shared and distinct transcriptomic cell types across neocortical areas. Nature 2018, 563 (7729), 72–78. DOI: 10.1038/s41586-018-0654-5 From NLM Medline.

(24) Kim, M. H.; Radaelli, C.; Thomsen, E. R.; Monet, D.; Chartrand, T.; Jorstad, N. L.; Mahoney, J. T.; Taormina, M. J.; Long, B.; Baker, K.; et al. Target cell-specific synaptic dynamics of excitatory to inhibitory neuron connections in supragranular layers of human neocortex. Elife 2023, 12. DOI: 10.7554/eLife.81863 From NLM Medline.

(25) Ooki, A.; Dinalankara, W.; Marchionni, L.; Tsay, J. J.; Goparaju, C.; Maleki, Z.; Rom, W. N.; Pass, H. I.; Hoque, M. O. Epigenetically regulated PAX6 drives cancer cells toward a stem-like state via GLI-SOX2 signaling axis in lung adenocarcinoma. Oncogene 2018, 37 (45), 5967–5981. DOI: 10.1038/s41388-018-0373-2 From NLM Medline.

(26) Krepela, E.; Vanickova, Z.; Hrabal, P.; Zubal, M.; Chmielova, B.; Balaziova, E.; Vymola, P.; Matrasova, I.; Busek, P.; Sedo, A. Regulation of Fibroblast Activation Protein by Transforming Growth Factor Beta-1 in Glioblastoma Microenvironment. Int J Mol Sci 2021, 22 (3). DOI: 10.3390/ijms22031046 From NLM Medline.

(27) Li, Y.; Xu, X.; Wang, X.; Zhang, C.; Hu, A.; Li, Y. MGST1 Expression Is Associated with Poor Prognosis, Enhancing the Wnt/beta-Catenin Pathway via Regulating AKT and Inhibiting Ferroptosis in Gastric Cancer. ACS Omega 2023, 8 (26), 23683–23694. DOI: 10.1021/acsomega.3c01782 From NLM PubMed-not-MEDLINE.

(28) Wess, M.; Andersen, M. K.; Midtbust, E.; Guillem, J. C. C.; Viset, T.; Storkersen, O.; Krossa, S.; Rye, M. B.; Tessem, M. B. Spatial integration of multi-omics data from serial sections using the novel Multi-Omics Imaging Integration Toolset. Gigascience 2025, 14. DOI: 10.1093/gigascience/giaf035 From NLM Medline.

(29) Tuck, M.; Grelard, F.; Blanc, L.; Desbenoit, N. MALDI-MSI Towards Multimodal Imaging: Challenges and Perspectives. Front Chem 2022, 10, 904688. DOI: 10.3389/fchem.2022.904688 From NLM PubMed-not-MEDLINE.

(30) Bowman, A. P.; Bogie, J. F. J.; Hendriks, J. J. A.; Haidar, M.; Belov, M.; Heeren, R. M. A.; Ellis, S. R. Evaluation of lipid coverage and high spatial resolution MALDI-imaging capabilities of oversampling combined with laser post-ionisation. Anal Bioanal Chem 2020, 412 (10), 2277–2289. DOI: 10.1007/s00216-019-02290-3 From NLM Medline.

(31) Ali, A.; Davidson, S.; Fraenkel, E.; Gilmore, I.; Hankemeier, T.; Kirwan, J. A.; Lane, A. N.; Lanekoff, I.; Larion, M.; McCall, L. I.; et al. Single cell metabolism: current and future trends. Metabolomics 2022, 18 (10), 77. DOI: 10.1007/s11306-022-01934-3 From NLM Medline.

(32) Kapalczynska, M.; Kolenda, T.; Przybyla, W.; Zajaczkowska, M.; Teresiak, A.; Filas, V.; Ibbs, M.; Blizniak, R.; Luczewski, L.; Lamperska, K. 2D and 3D cell cultures - a comparison of different types of cancer cell cultures. Arch Med Sci 2018, 14 (4), 910–919. DOI: 10.5114/aoms.2016.63743 From NLM PubMed-not-MEDLINE.

(33) Rood, J. E.; Wynne, S.; Robson, L.; Hupalowska, A.; Randell, J.; Teichmann, S. A.; Regev, A. The Human Cell Atlas from a cell census to a unified foundation model. Nature 2025, 637 (8048), 1065–1071. DOI: 10.1038/s41586-024-08338-4 From NLM Medline.

(34) Wang, L.; Jung, J.; Babikir, H.; Shamardani, K.; Jain, S.; Feng, X.; Gupta, N.; Rosi, S.; Chang, S.; Raleigh, D.; et al. A single-cell atlas of glioblastoma evolution under therapy reveals cell-intrinsic and cell-extrinsic therapeutic targets. Nat Cancer 2022, 3 (12), 1534–1552. DOI: 10.1038/s43018-022-00475-x From NLM Medline.

(35) Scupakova, K.; Dewez, F.; Walch, A. K.; Heeren, R. M. A.; Balluff, B. Morphometric Cell Classification for Single-Cell MALDI-Mass Spectrometry Imaging. Angew Chem Int Ed Engl 2020, 59 (40), 17447–17450. DOI: 10.1002/anie.202007315 From NLM Medline.

