## Supplemental figures MALDI-MSI and SPT for "One Section, Two Worlds: Single-Cell Integration of MALDI-MSI and Spatial Transcriptomics on the Same Single Tissue Section"

**Supplementary table 1: List of gene markers per cell type used for cell identification and identified cells.**

| **Cell type** | **Gene markers** | **Identified cells** |
|---|---|---|
| Astrocytes | AQP4, GJA1, FGFR3 | 2009 |
| Endothelial cells | PECAM1, FLT1, NRP1 | 530 |
| Excitatory neurons | CRYM, SLC17A7, SLC17A6 | 253 |
| GSCs | PAX6, SOX2 | 594 |
| Inhibitory neurons | GAD1, SST, WIF1 | 189 |
| MES | TGBF1, MGST1, NCSTN | 365 |
| Oligodendrocytes | CAPN3, MOBP, MOG, OPALIN | 2174 |
| OPC | PDGFRA, SOX10, OLIG2 | 653 |
| T-cells | CD4 | 686 |
| TAMs | CD163, CD68, P2RY12, CX3CR1, TREM2 | 865 |

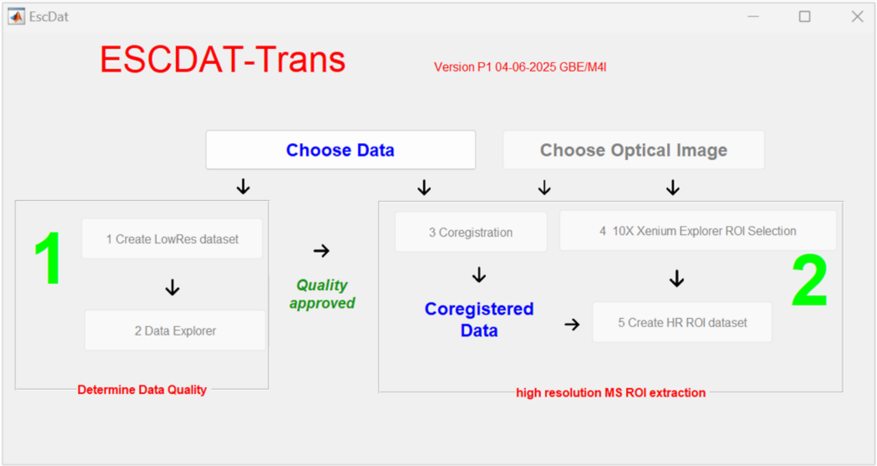

| GUI Button | Scriptname | Script function |
|---|---|---|
| Choose data | EscDat.m pushbutton8_Callback | Choose TimsTOF .d data folder |
| Choose optical image | EscDat.m pushbutton7_Callback | Choose microscope image |
| 1. Create LowRes dataset | ReadShowDataLowRes.m → Read_my_tsf_data_LRdataset.py | Read BRUKER binary TIMS-TOF tsf data using Python Create MATLAB 1 Da low resolution MS image |
| 2. Data Explorer | HistDistGUIsparce.m | Explore MATLAB 1 Da low resolution MS image and create TIC, single mass channel or combined mass channels image for coregistration with µscoop (fluorescense, stained) image B |
| 3. Coregistration | ECoRegT. m → Coregister.m | Coregister (and rotate/mirror if needed) A and B Store registration points and tranformation matrix |
| 4. 10X Xenium Explorer Selection | GeoJSON creation from 10X software | Run 10X or another package to analyze image A (annotate cells end nuclei), export Object data (All Objects) as GeoJSON |
| 5. Create HR ROI dataset | CreateHDRDataROI.m | Load tranformation matrix<br>Load geoJSON file from 10X<br>Load mass image mim: imcounts, Xpos, Ypos<br>Load mass image (mim) rotate/flip info mimmod<br>Flag for cell or nucleus:<br>UseNuc=2 Cell with nucleus<br>Output: text file ROIsOn.txt with structure: %d %d %s %d : i n name group<br>where i = index number of cell in geoJSON file, cells without nucleus are skipped n = number of overlapping pixels name = Cell identifier group = Group/category number<br>%8.6f = n times weighing factor of pixel<br>%d = n times X coordinate of pixel<br>%d = n times Y coordinate of pixel |
| | PlotTestRead.py | Read output from CreateHDRDataROI.m Extract from BRUKER binary TIMS-TOF tsf data using Python and sum all X Y spectra and weigh by weight factor Output: text file with 2 rows: m/z value Intensity ROIsSummedSpectrum.txt |
| | peapi EscDat version | read ROIsSummedSpectrum.txt<br>Perform peakpicking (see Eijkel et. al. Surface and Interface Analysis Volume 41, Issue 8, Pages 675 - 685August 2009 )<br>output matlab peaklist |
| | Create PeakPickedSpectra.py | Use peaklist to create HR ms data from GeoJSON annotations Extract from BRUKER binary TIMS-TOF tsf data using Python and sum all X Y spectra and weigh by weight factor Output: CSV file with cell-id from GeoJSON, all GeoJSON metrics and the pixel weighed spectrum for that cell with/without nucleus or nucleus CSV file with Seurat formatted IMS data |

**Supplementary figure 1: ESCDAT Software GUI and explanation.**

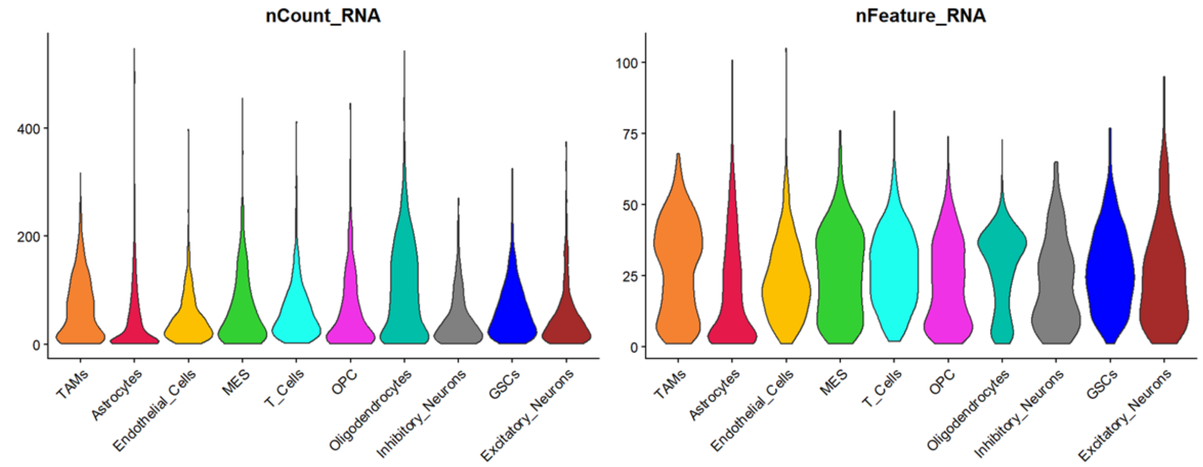

**Supplementary figure 2: Distribution of gene counts and gene features per cell type.** Violin plots of total UMI counts per cell (nCount_RNA) and number of detected genes per cell (nFeature_RNA) are shown for each predicted cell-type cluster; TAMs, astrocytes, endothelial cells, MES, t-cells, OPC, oligodendrocytes, inhibitory neurons, GSCs, excitatory neurons. Comparable distributions across all groups demonstrate uniform sequencing depth and gene-detection efficiency, with no cluster-specific biases affecting downstream analyses.

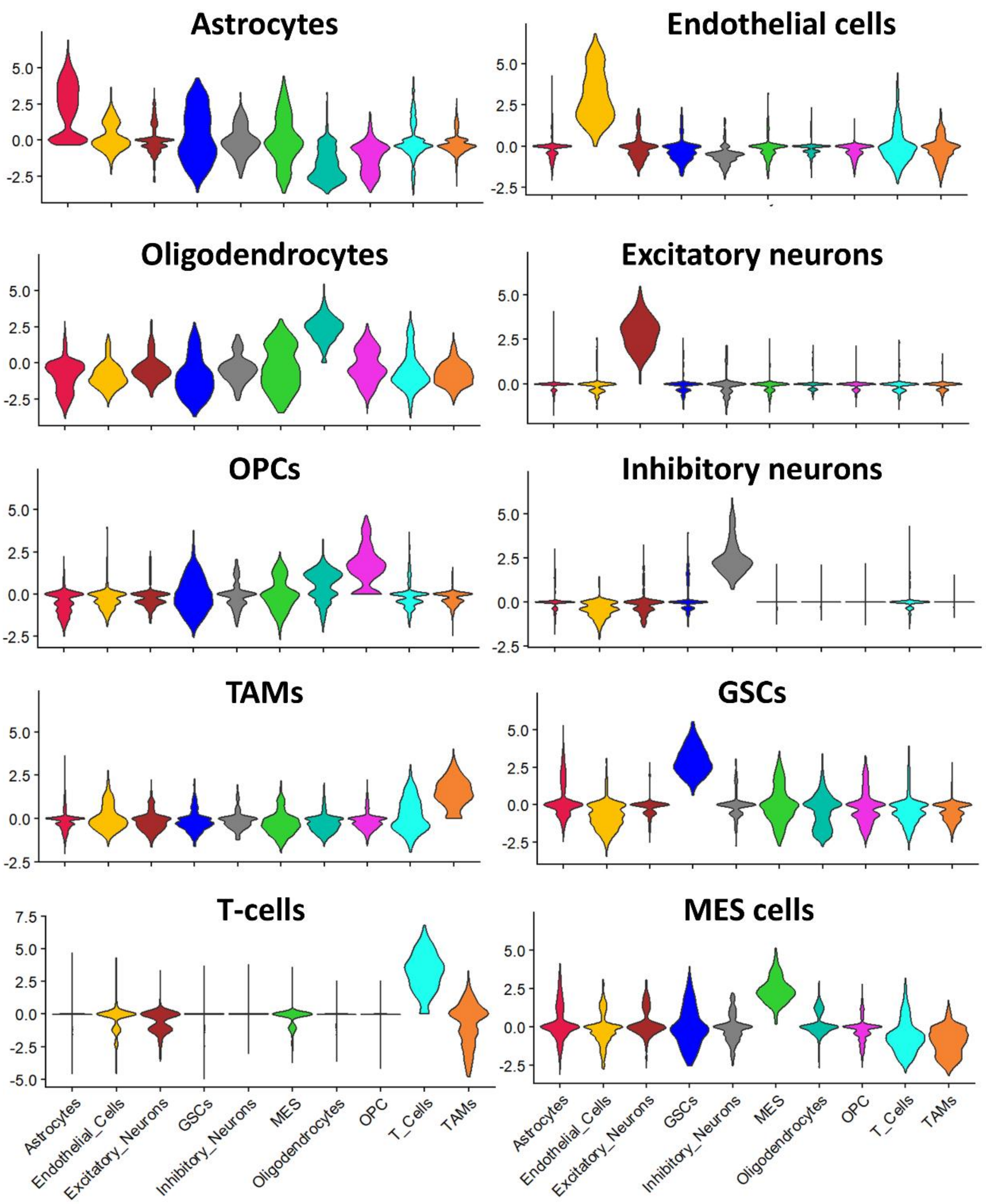

**Supplementary figure 3: Score distribution per cell type.** For each of the ten selected cell-type signatures we calculated an AddModuleScore per cell and then assigned each cell to the signature with its highest score. In every violin plot, the y-axis shows the range of module scores for that signature, and the x-axis groups cells by their assigned cell type. The signature score is sharply enriched in its matching cell-type cohort while remaining near zero in all other groups. However, low background signatures can be seen in cell types, possibly reflecting baseline expression.

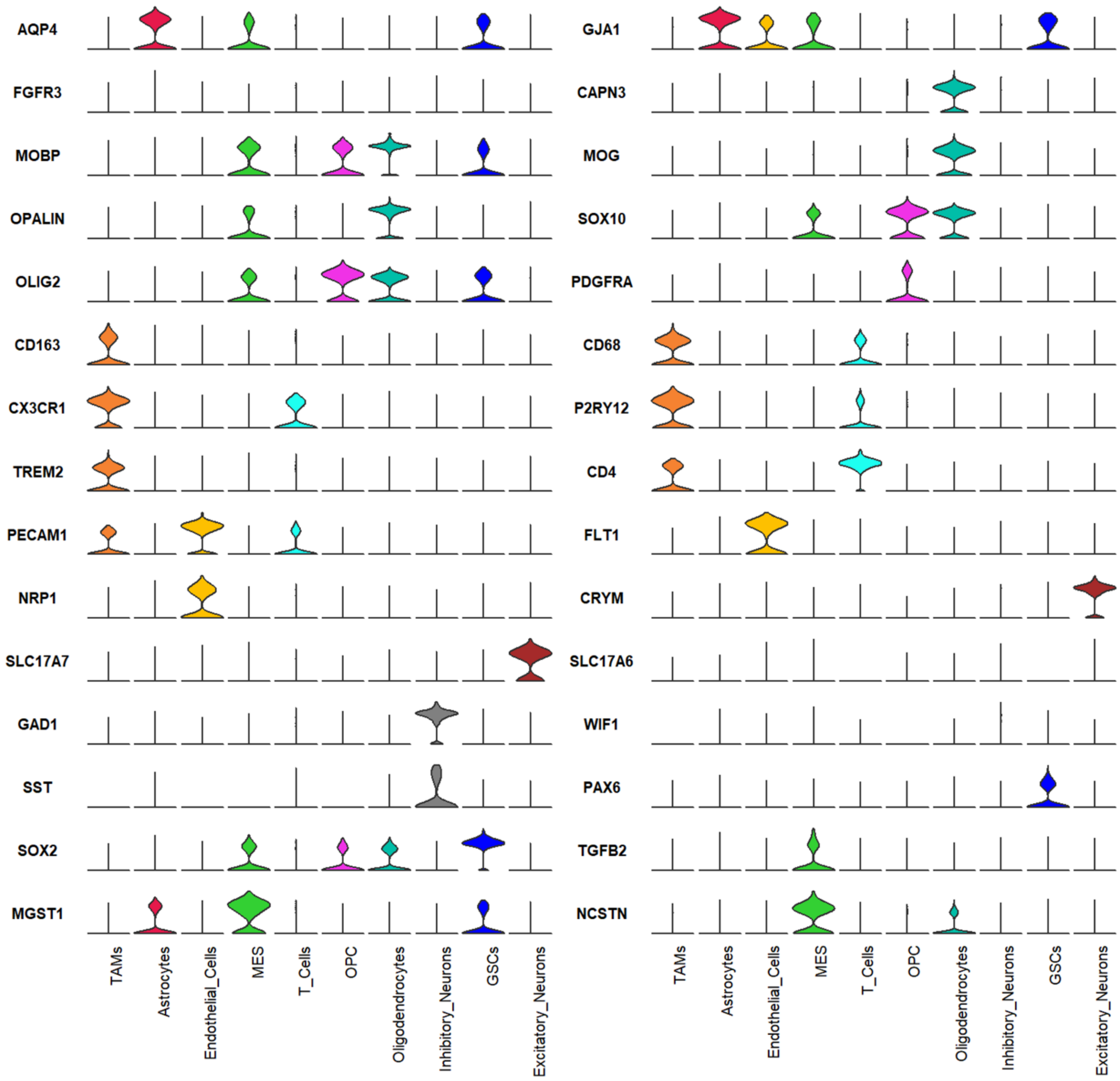

**Supplementary figure 4: Score distribution of individual marker-gene specificity across predicted cell types.** For each marker gene, normalized Xenium RNA expression is shown as a violin density plot across all cells, grouped by their predicted cell type.

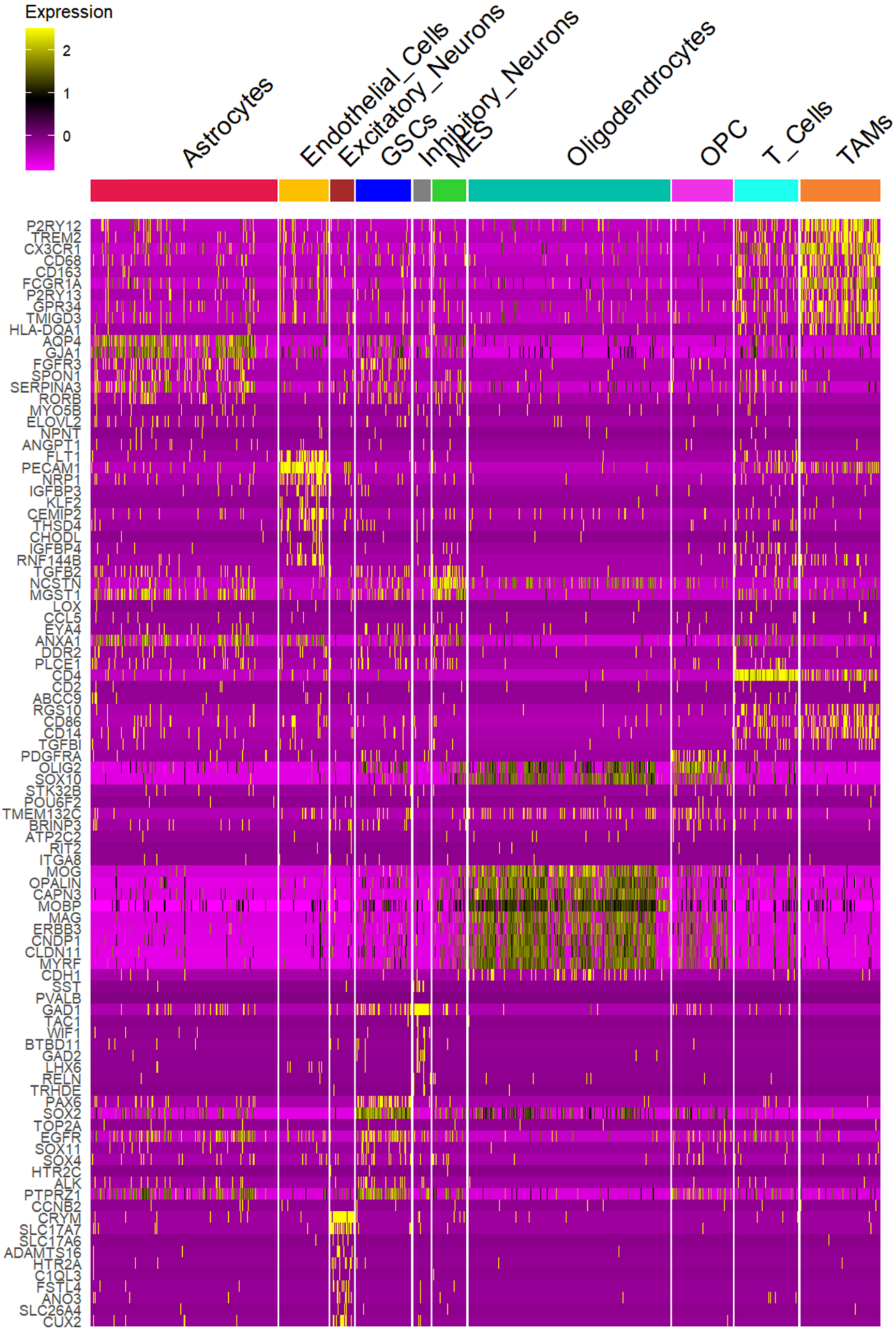

**Supplementary figure 5: Heatmap of top RNA markers across predicted cell types.** Normalized RNA expression of the top ten positive marker genes for each predicted cell-type cluster identified via Seurat displayed in a heatmap.

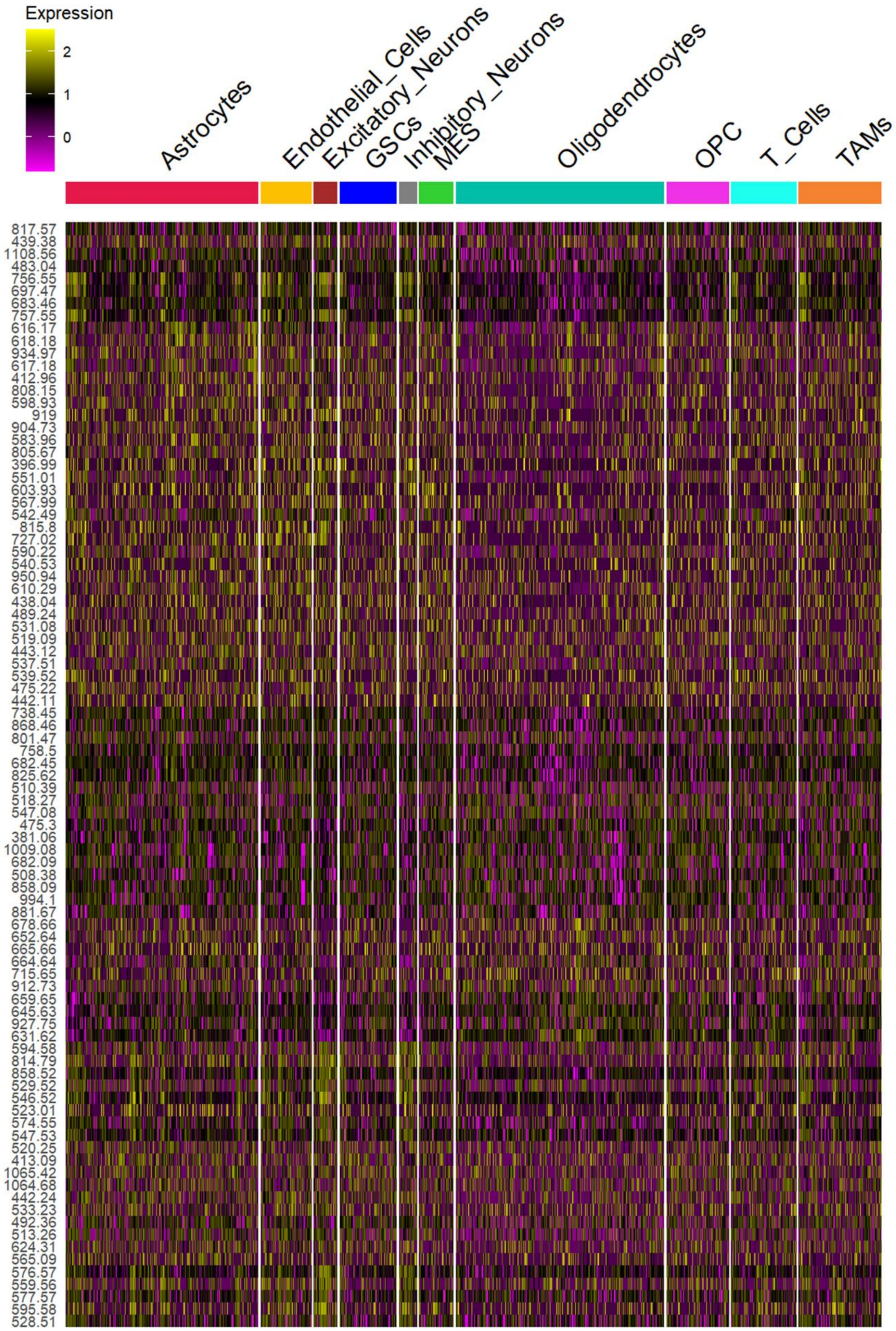

**Supplementary figure 6: Heatmap of top m/z markers across predicted cell types.** Normalized *m/z* expression of the top ten positive marker *m/z* values for each predicted cell-type cluster identified via Seurat displayed in a heatmap.

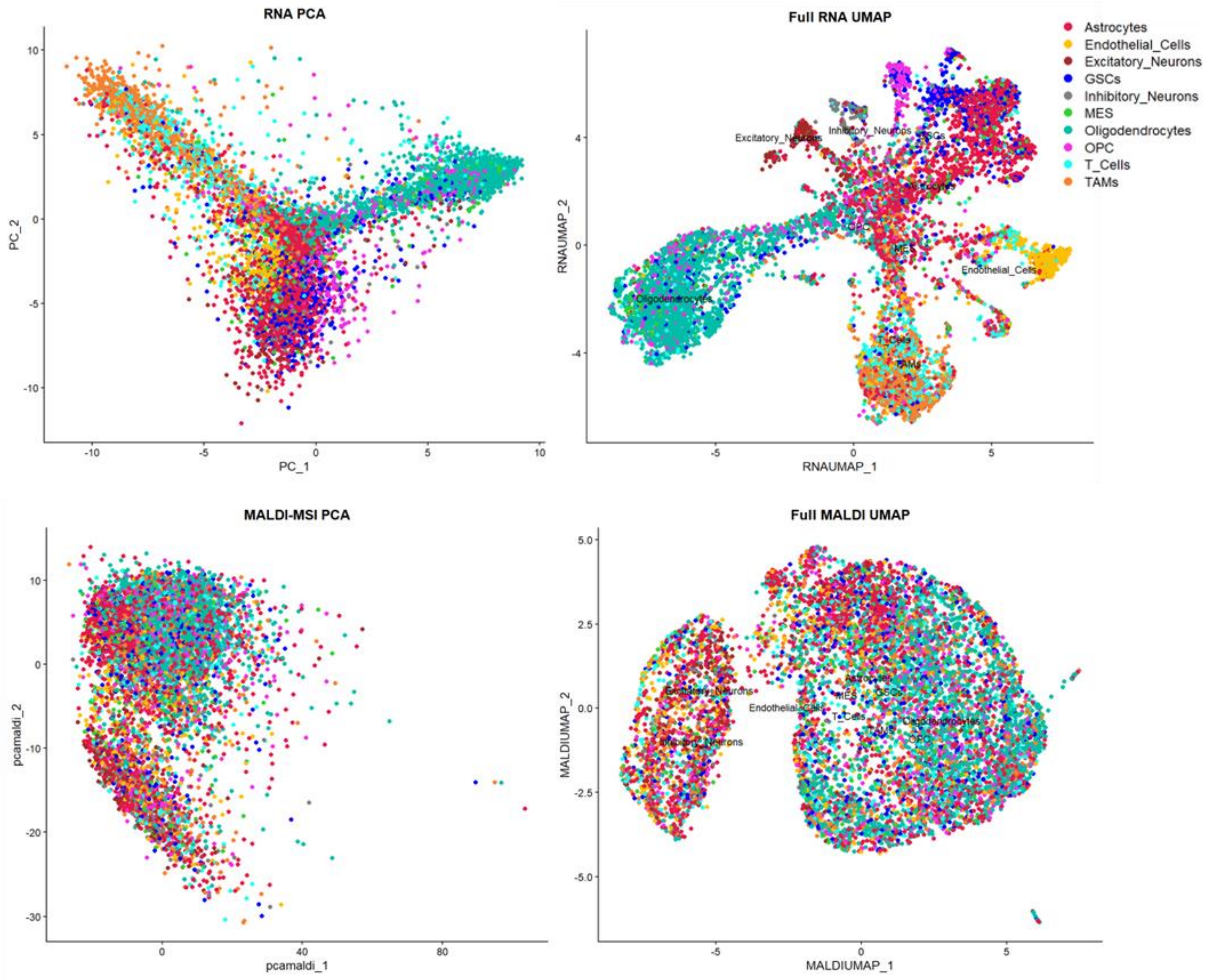

**Supplementary figure 7: Modality-specific dimensionality reduction.** (A) RNA PCA (left) and RNA UMAP (right) embeddings of n = 8,318 cells based on normalized spatial transcriptomic data. Distinct clusters corresponding to major cell types are annotated. (B) MALDI-MSI PCA (left) and MALDI-MSI UMAP (right) embeddings of the same cells based on TIC-normalized mass spectrometry intensities. Cell types intermix, illustrating the lower resolution of MALDI-MSI alone.

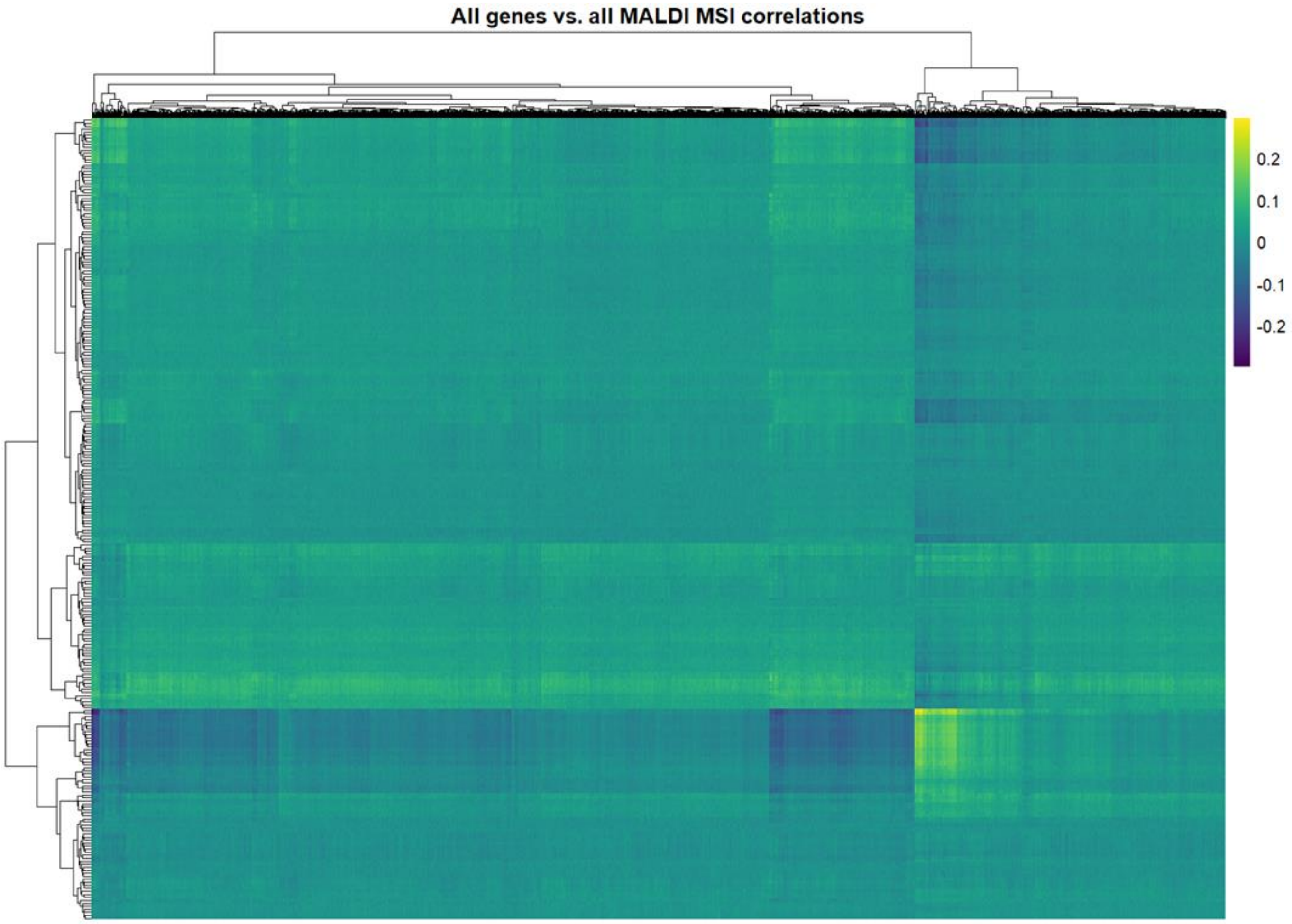

**Supplementary Figure 8. Global gene – MALDI-MSI correlation heatmap.** Pearson correlation coefficients between normalized expression of all genes (horizontal) and intensities of all MALDI-MSI *m/z* features (vertical) across all single-cell profiles. Both rows and columns are hierarchically clustered (dendrograms at top and left) to reveal co-varying modules. This map shows distinct gene-metabolite/lipid clusters.
